# Amyloid Beta Pathology Accelerates Alterations in the Visual Pathway of the 5xFAD Mouse

**DOI:** 10.1101/2024.09.29.615699

**Authors:** Shaylah McCool, Jennie C. Smith, Asia Sladek, Shan Fan, Matthew J. Van Hook

## Abstract

Alzheimer’s disease (AD) is a neurodegenerative disorder characterized by the formation of amyloid beta plaques and neurofibrillary tangles that leads to decreased quality of life due to behavioral, motor, and cognitive impairments. Due to the widespread pathological nature of AD, many brain regions are affected by amyloid beta plaques including regions important for vision such as the lateral geniculate nucleus (LGN) of the thalamus which is critical for relaying signals from the retina to the primary visual cortex. Using a wide range of techniques including electrophysiological approaches, *in vivo* and *ex vivo* imaging methods, and immunohistochemistry in a mouse model with progressing amyloidosis (5xFAD), the goal of this study was to determine whether AD-like pathology disrupts neuronal and synaptic structure and function in the visual system. *In vivo* electroretinogram recordings revealed photoreceptor dysfunction in the 6- and 9-month-old 5xFAD mice, while optical coherence tomography indicated no changes in retinal thickness. In the dorsolateral geniculate nucleus (dLGN), the rodent homolog of the primate LGN, we identified decreased densities of retinal ganglion cell axon terminals and fewer thalamocortical (TC) neuron cell bodies. No detectable deficits in excitatory synaptic function or TC neuron dendritic structure were seen in the dLGN, and reflexive visual behavior was also found to be normal in the 5xFAD mice. These results indicate relatively modest amyloid-triggered dysfunction in these stages of the visual system suggesting that amyloid beta plaque formation may play only a small role in the visual system dysfunction seen in AD patients. These results may also point to potential compensatory mechanisms that preserve function of visual pathways in the 5xFAD visual system.

## Introduction

Alzheimer’s disease (AD) is a leading cause of death worldwide, one that is rising in prevalence with the aging population. This neurodegenerative disorder is defined by a loss of memory and cognitive impairment that leads to behavioral and motor deficits due to a myriad of risk factors including age, genetics, and environment [1,2]. The pathological progression of AD involves formation of amyloid beta (Aβ) plaques preceding and inducing formation of neurofibrillary tangles (NFTs) caused by hyperphosphorylated tau followed by cerebral atrophy [3]. Individuals with AD experience visual manifestations of the disease as some of the first reported symptoms with untreated vision loss a potential modifiable risk factor to slow progression of the disease as studies report a bidirectional relationship between visual impairments and cognitive impairments and dementia [4-7]. These visual manifestations of AD include decreases in visual acuity, contrast sensitivity, and color vision in addition to visual field loss, ocular motor dysfunction, and subpar visual attention [8,9]. Pathological changes found in post-mortem AD patients that may contribute to these visual impairments include reduced retinal thickness and vasculature, fewer optic nerve axons, and Aβ plaques and NFTs in both cortical and subcortical regions of the brain important for vision [8]. Whether the visual deficits in AD patients are due to Aβ plaques, NFTs, or some other mechanistic change is unknown.

Currently, physicians must rely on cognitive testing, imaging methods, and CSF or blood tests for identification of AD biomarkers at which point the disease has progressed to the point where decline in quality of life is inevitable. Early detection is a critical component of AD research, and the visual system has great potential for early detection due to its accessibility and the ability to test many aspects of vision clinically.

The focus of this study was to determine the impact of AD-like amyloid pathology on the structure and function of neurons and synapses in the retina and the dLGN, a retinorecipient region of the thalamus that is critical for relaying information from the retina to the cortex for conscious vision. Working under the hypothesis that Aβ pathology leads to functional and structural disruption of the neurons and synapses in visual pathways, we sought to identify mechanisms underlying visual deficits in AD to provide insight on the effects of Aβ plaques on the visual system using a wide array of techniques to test the overall health of the retinogeniculate pathway.

Overall, our data point to disruptions in the function of photoreceptors without apparent retinal degeneration in mice up to 12mo of age. Studies of dLGN structure and function indicate a modest loss of retinal ganglion cell axon terminals and of postsynaptic dLGN neurons without detectable effects on synaptic function of surviving neurons. Finally, measures of optomotor responses point to intact reflexive visual behavior. These results indicate that amyloid pathology has fairly modest effects on visual pathway structure and function in the 5xFAD mice suggesting that other elements of AD-like pathology might be significant contributors of visual manifestations of AD.

## Methods and Methods

### Animals

The Institutional Animal Care and Use Committee at the University of Nebraska Medical Center approved all animal protocols. Both male and female 5xFAD mice on a C57BL/6J background and C57BL/6J control mice were purchased from Jackson Laboratories [B6.Cg-Tg(APPSwFlLon,PSEN1*M146L*L286V)6799Vas/Mmjax (5XFAD, The Jackson Laboratory #034848-JAX, RRID:MMRRC_034848-JAX) and C57BL/6J (B6, The Jackson Laboratory #:000664, RRID:IMSR_JAX:000664)] and housed on a 12 h light/dark cycle except for overnight dark-adaptation for ERG recordings with access to food and water *ad libitum*. The 5xFAD mice do not contain the retinal degeneration allele Pde6b^rd1^. Animals were killed via inhalation of CO_2_ and cervical dislocation.

### Histology & Immunofluorescence Staining

After euthanasia by CO_2_ inhalation and cervical dislocation, the brain was quickly dissected, washed with 1x PBS, and fixed in 4% PFA for 4 hrs. The brains were cryoprotected in 30% sucrose in 1x PBS and later embedded in 3% agar. A Leica VT1000S vibratome was used to slice 50 μm sections of tissue, which were then mounted on Superfrost Plus slides (Fisher) for staining.

Thioflavin S was used to stain brain tissue to visualize Aβ plaques in the dLGN. Brain slices mounted on slides were washed with 70% ethanol for 1 min followed by a 1 min wash in 80% ethanol. Slides were incubated in a filtered solution of 1% thioflavin S in 80% ethanol for 15 min after which they were washed in an 80% ethanol solution for 1 min, a 70% ethanol solution for 1 min, and ending with two 1 min washes in distilled water. Coverslips were mounted using Vectashield Hardset mounting medium, and slides were allowed to dry in the dark. Slides were stored at 4°C until imaging. Sections of dLGN tissue were imaged at 10x using a fixed-stage upright microscope (Olympus BX51-WI) with 20 ms exposure. For analysis of Aβ density, ImageJ was used to measure the dLGN area, and the Cell Counter plug-in was used to determine plaque density in each slice.

For vGlut2 and NeuN immunofluorescence staining, slides were rinsed in 1x PBS and blocked for 1 hr followed by an overnight incubation at 4°C in a 1:500 primary antibody solution (guinea pig anti-vGlut2, Millipore Cat# AB2251-I, RRID:AB_2665454) or (guinea pig anti-NeuN, Millipore Cat# ABN90, RRID:AB_11205592). The slides were washed 6x10 min in 1x PBS and blocked for 1 hr. A 2 hr incubation in a 1:250 secondary antibody solution (for vGlut2: goat anti-guinea pig 488, Thermo Fisher Scientific Cat# A-11073, RRID:AB_2534117) or (for NeuN: goat anti-guinea pig 568, Thermo Fisher Scientific Cat# A-11075, RRID:AB_2534119) at room temperature was followed by three 10 min washes in 1x PBS followed by a 1 min wash in dH_2_O. Vectashield Hardset mounting medium was used to coverslip the slides which were then stored at 4°C until imaging. A 2-photon microscope with a Ti:sapphire laser tuned to 800 nm was used to image the tissue with a 185 μm x 185 μm field of view for vGlut2 and a 370 μm x 370 μm field of view for NeuN. The Synapse Counter plug-in was used to detect vGlut2 puncta in ImageJ from a single section of the tissue (Smith, 2022). The Cell Counter plug-in was used to count the presence of NeuN-labeled cell bodies in a 40-μm-thick section of the tissue.

### Electroretinogram (ERG)

For ERG recording, mice were dark-adapted overnight (>12 hrs) and anesthetized with an intraperitoneal ketamine/xylazine injection [100 mg/kg ketamine (Zoetis, Parsipanny, NJ) and 5 mg/kg xylazine (Akorn Inc., Lake Forest, IL)]. A drop of tropicamide (Akorn Inc., Lake Forest, IL) was applied to each eye followed by a drop of proparacaine (Alcon Laboratories, Fort Worth, TX). Both eyes received a generous amount of 0.3% hypromellose gel (GenTeal Tears, Alcon, Fort Worth, TX) after which the animals were positioned on the Celeris Diagnosys small animal ERG system, and an electrode/fiber optic stimulator was placed on both eyes. Scotopic ERGs were recorded in response to stimulus intensities of 0.0005, 0.001, 0.005, 0.01, 0.05, 0.1, 0.5, 1, and 3 cd*s/m^2^. ERGs were bandpass filtered during acquisition (0.125 to 300 Hz) and sampled at 2kHz. Five responses were averaged for 0.0005 – 0.01 cd*s/m^2^ stimuli (10 s inter-flash interval) and three responses were averaged for 0.05 and 0.1 cd*s/m^2^ (17 s inter-flash interval) and for 0.5 – 3 cd*s/m^2^ (22 s inter-flash intervals). Analysis of A- and B-wave amplitudes was performed using Clampfit 11.2 (Molecular Devices, Sunnyvale, CA). A-waves were measured from baseline to the negative A-wave trough, while B-waves were measured from the A-wave to the rounded top of the B-wave. Oscillatory potentials (OPs) were isolated off-line by band-pass filtering between 65 and 280 Hz, and amplitudes were measured from baseline. Four OPs per trace were summed for our analysis.

### Optical Coherence Tomography (OCT)

Mice were anesthetized with an intraperitoneal ketamine/xylazine injection. Tropicamide and proparacaine were applied to each eye followed by a 0.3% hypromellose gel (GenTeal Tears, Alcon, Fort Worth, TX). A 3.2 mm contact lens (base curve 1.70, plano power) was placed on the eye prior to imaging. Heidelberg Spectralis OCT machine (Heidelberg Engineering, Inc., Franklin, MA, United States) adapted with a +35 diopter lens was used to acquire cross-sectional images of the retina at 12 axes. Analysis was conducted with Heidelberg OCT software. The retinal nerve fiber layer (RNFL), ganglion cell layer (GCL), and inner plexiform layer (IPL) thickness was measured as a group. The outer nuclear layer (ONL) and external limiting membrane (ELM) thickness was measured as a group. Using three images, analysis was performed by two graders after which a third grader was employed if measurements between the first two graders had a discrepancy larger than 10 μm between values. If all three graders’ measurements were different by more than 10 μm, that section was excluded from analysis. The values between graders were averaged resulting in three measurements per retinal layer per eye. A nested one-way ANOVA was performed using these measurements.

### Patch clamp electrophysiology

Mice were killed via inhalation of CO_2_ and cervical dislocation followed by decapitation and extraction of the brain into cold artificial cerebrospinal fluid (aCSF: 128 mM NaCl, 2.5 mM KCl, 1.25 mM NaH_2_PO_4_, 24 mM NaHCO_3_, 12.5 mM glucose, 2 mM CaCl_2_, and 2 mM MgSO_4_; 300 – 315 mOsm with a pH of 7.4 continuously bubbled with 5% CO_2_ / 95% O_2_). For slice preparation, we used the “protected recovery” method [10,11]. The brain was removed from the aCSF after 60 sec, and the cerebellum was removed to create a flat surface which was then glued to the stage of the Leica VT 1000s vibratome in front of an agar block for support. Coronal slices, 250-μm-thick, containing the dLGN, were created while submerged in ice-cold aCSF bubbled with 5% CO_2_ / 95% O_2_. The slices were hemisected and placed into a 32°C bubbling N-methyl-D-glucamine (NMDG) solution for 12 – 20 min after which they were moved to bubbling, room temperature aCSF for 1 hr to recover from slicing. During data collection, the slices were superfused with aCSF in a recording chamber on a fixed-stage upright microscope (Olympus BX51-WI).

For recording miniature excitatory post-synaptic currents (mEPSCs), we used an intracellular pipette solution of Cs-methanesulfonate (120 mM Cs-methanesulfonate, 2 mM EGTA, 10 mM HEPES, 8 mM TEA-Cl, 5 mM ATP-Mg, 0.5 mM GTP-Na_2_, 5 mM phosphocreatine-Na_2_) supplemented with 2mM QX314. Intracellular solution pH was adjusted to 7.4 and osmolarity was adjusted to 270 – 275 mOsm. Pipettes were pulled to have ∼ 1 μM openings and a resistance of ∼ 4 – 7 MOhms when filled with the intracellular pipette solution. The aCSF was supplemented with 60 μM picrotoxin during mEPSC recording. Within the dLGN core, we targeted thalamocortical (TC) neurons by shape and size of soma. Whole-cell recordings were conducted, and TC neuron identity was confirmed by the presence of a low-voltage activated calcium current. A holding potential of -70 mV (after correcting for liquid junction potential) was used to record miniature excitatory postsynaptic currents. To analyze the mEPSC traces, they were first filtered with a lowpass filter at 500 Hz in ClampFit 11.2. Using MiniAnalysis software (Synaptosoft, Fort Lee, NJ, USA) and approximately 100 mEPSCs were analyzed from each cell following detection using the following parameters: threshold = 4 pA, period for local max = 10000 μs, time for baseline = 10000 μs, period for decay time = 10000 μs, fraction to find decay time = 0.37, period to average for baseline = 5000 μs, area threshold = 6 pC, points to average for peak = 3, direction = negative.

For neurobiotin cell fills and dendritic reconstructions from a separate set of recordings, the Cs-methanesulfonate pipette solution was supplemented with 2% neurobiotin and 2% CF568 dye and TC neurons were targeted for whole cell patch clamp recording. To fill the cell with the dye, we either applied a 500 pA square wave pulse (+/- 250 pA) at 2 Hz or allowed the cell to fill via passive diffusion. The slice was then transferred to a well plate where it was submerged in 4% paraformaldehyde for 1 hr. After fixation and 3x10 min washes with 1x PBS, the slice was incubated in a solution of 1x PBS, 20% Triton X-100, and streptavidin-568 while rocking at 4°C. After 1 week of incubation, the slice was rinsed 3x10 min in 1x PBS and transferred to a slide and coverslipped using Vectashield Hardset prior to imaging. The filled cells were imaged using a 2-photon microscope with a Ti:sapphire laser tuned to 800 nm. ImageJ Simple Neurite Tracer plugin was used to reconstruct the dye-filled neuron after which a Sholl analysis was performed to determine the number of dendritic crossings per 10 μm radius from the center of the cell.

### Optomotor Response (OMR)

The custom OMR apparatus consists of four monitors surrounding a raised pedestal where the mouse is placed during recording. The camera recording the movements of the mouse is directly above the pedestal. The floor of the set-up is a mirror which reflects the stimuli presented on the monitors. A baseline recording is acquired followed by a 5 min acclimation period in which the mouse sits on the pedestal prior to testing. To test visual acuity, a stimulus was used that consists of vertical lines ranging in spatial acuity (0.025 to 0.450 cpd). To test contrast sensitivity, a stimulus was used that consists of vertical lines ranging in contrast (1 - 96% contrast). These stimuli are presented on the monitors and rotate clockwise and counterclockwise. Analysis was blinded and performed manually.

### Statistical Analyses

Statistical analyses were performed using GraphPad Prism 10 software. Two-way ANOVA with Tukey’s multiple comparisons was used to analyze immunohistochemistry data, electroretinograms, cell fills, and optomotor response data. For analysis of optical coherence tomography and miniature excitatory postsynaptic current recordings, a nested one-way ANOVA with Tukey’s multiple comparisons was used. We tested mEPSC frequency for a normal distribution using the D’Agostino-Pearson normality test. Non-normally distributed data were log transformed prior to nested statistical analyses. Nested analyses are reported as number of eyes or cells as individual data points. Two-way ANOVAs are reported as individual data points with means ± SEM. Sample sizes are reported in figure legends.

## Results

To test whether retinal function is altered in mice with amyloid pathology, we performed dark-adapted electroretinogram (ERG) recordings on 6-, 9-, and 12-month 5xFAD and age-matched C57 mice (Fig. 1.A). At the highest stimulus intensity (3 cd*s/m^2^) in 6-month-old 5xFAD mice, we found an increase in both A- and B-wave amplitudes compared to age-matched controls (p=0.0144 and p=0.0315, respectively). However, when we recorded ERGs at the 9-month time point, we found lower A- and B-wave amplitudes in the 5xFAD mice compared to controls (p=0.0008 and p=0.0041, respectively). At 12 months of age, we observed no difference in ERG wave amplitudes between genotypes (A-wave p=0.7931 and B-wave p=0.4283). Considering the C57 mice at all three timepoints, we found a significant decrease in ERG wave amplitudes between 9 months and 12 months (A-wave p=0.0075 and B-wave p=0.0029) pointing to an age-dependent decline in ERG responses consistent with prior reports [12,13]. With the 5xFAD mice, however, we saw a significant decrease in ERG wave amplitudes between 6 months and 9 months (both p<0.0001) indicating an age-related acceleration in decline of retinal function in our 5xFAD mice (Fig. 1.B and 1.C). As bipolar cells are postsynaptic to photoreceptors, we next sought to test whether the reduced B-wave was the result of dysfunction at the photoreceptor-to-bipolar cell synapse or a consequence of reduced responsiveness of photoreceptors. Taking the ratio of the B-wave to A-wave amplitude, we were able to assess synaptic function finding no significant difference between our groups which points to normal synaptic transmission between the photoreceptors and bipolar cells (Fig. 1.D) and indicating that retinal dysfunction seen via ERG recordings originates with the photoreceptors. In a study of human retina, the AD retina showed altered proteomic analysis in photoreceptor-related pathways [14].

**Figure 1.**
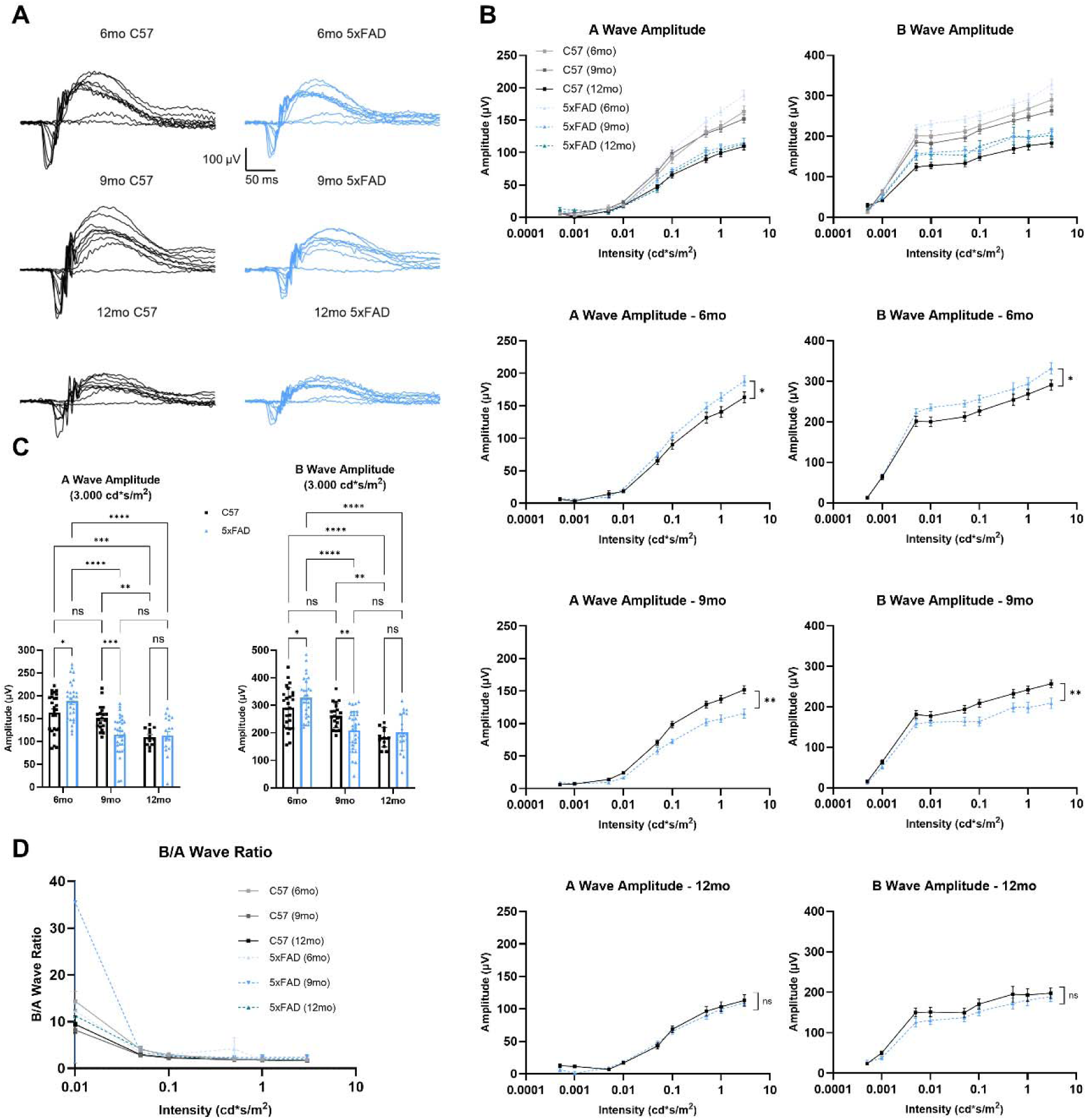
Electroretinogram (ERG) recordings from 6-, 9-, and 12-month mice show deficits in 5xFAD photoreceptors. (A) Example ERG traces from C57 (black) and 5xFAD (blue) mice at 3 timepoints show waveforms produced by 9 stimulus intensities ranging from 0.0005 – 3.0000 cd*s/m^2^. (B) Significantly altered A- and B-wave amplitudes are seen in the 6-month and 9-month 5xFAD mice. Two-way ANOVA with Tukey’s multiple comparisons post-hoc test was used for analysis. (C) Quantification of A- and B-wave amplitudes at the highest intensity stimulus (3.0000 cd*s/m^2^) show the significant increase and decrease in retinal function in the 6- and 9-month 5xFAD mice, respectively, analyzed by two-way ANOVA with Tukey’s multiple comparisons. (D) Ratio of B-wave to A-wave shows no significant difference between groups indicating normal synaptic function as analyzed by two-way ANOVA and Tukey’s multiple comparisons test. Sample size described as number of eyes: n = 26 (6mo C57), 30 (6mo 5xFAD), 20 (9mo C57), 34 (9mo 5xFAD), 12 (12mo C57), 20 (12mo 5xFAD).

To gain insight into the function of the inner retina, we analyzed ERG oscillatory potentials (OPs) (Fig. 2.A). These higher frequency, lower amplitude wavelets are found on the rising phase of the B-wave and originate from amacrine cells in the inner retina [15,16]. We found a pattern of OP dysfunction that mirrored the effects seen in A- and B-wave recordings (Fig. 2.B). Specifically, we found a significant increase in the sum of OP amplitudes in the 6mo 5xFAD mice compared to controls (p=0.0218). By 9 months, there was a significant decrease in the sum of OP amplitudes in 5xFAD mice compared to controls. At 12 months, there was no difference in these amplitudes between C57 and 5xFAD mice (p=0.0038 and p=0.4099, respectively). Again, there was a significant decrease in the sum of OP amplitudes of C57 mice between 9 and 12 months, while we found this decrease in amplitudes occurring between 6 and 9 months in the 5xFAD mice. These results point to dysfunction of photoreceptors in the 6- and 9-month 5xFAD retinas with the altered signal being carried to downstream components of the retinal circuit.

**Figure 2.**
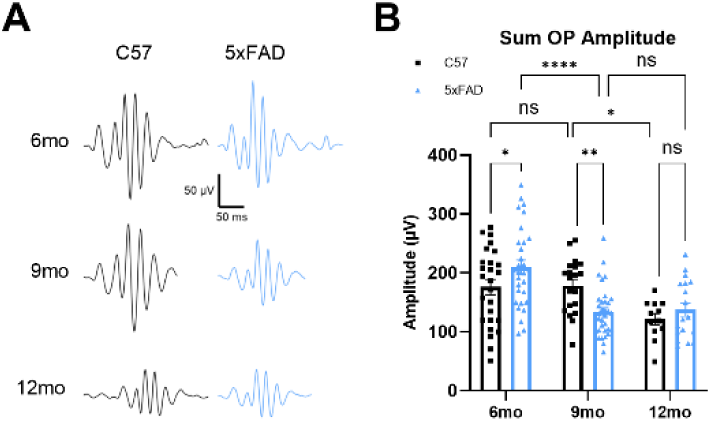
Oscillatory potential (OP) analysis of 6-, 9-, and 12-month ERGs show continued dysfunction in the inner retina reflecting A- and B-wave results. (A) Example OP traces from C57 (black) and 5xFAD (blue) mice at 3 timepoints show significantly increased OP amplitudes in the 6-month 5xFAD mice and significantly decreased OP amplitudes in the 9-month 5xFAD mice. (B) Quantification of the sum of the OP amplitudes shows significantly increased and decreased OP amplitudes in the 6- and 9-month 5xFAD mice, respectively, followed by equivalently decreased OP amplitudes by 12 months between genotypes similar to the results in Figure 1. Sample size described as number of eyes: n = 26 (6mo C57), 30 (6mo 5xFAD), 20 (9mo C57), 34 (9mo 5xFAD), 12 (12mo C57), 19 (12mo 5xFAD).

With our ERG results indicating dysfunction in the retina, we next aimed to investigate whether this is apparent as a parallel disruption of retinal structure. Using optical coherence tomography (OCT), a noninvasive imaging technique, cross-sectional images of the retina were collected, and layer thickness measurements were analyzed allowing us to determine whether there are alterations in thickness of the inner and outer retina (Fig. 3.A). There are conflicting studies on the topic of OCT layer thickness in 5xFAD mice. Some studies show that the inner and outer retina have reduced thickness in the 5xFAD mice [17], while others show the RNFL of 5xFAD mice is thinner at both younger and older timepoints with the IPL being thicker at 6mo [18]. To assess photoreceptor structural integrity, we measured the combined thickness of the external limiting membrane (ELM) and the outer nuclear layer (ONL), which contains the photoreceptor cell bodies. We saw no change in outer retinal thickness at either 500 or 1000 μm from the optic nerve head (Fig. 3.B). To assess inner retinal structure, we measured the combined thickness of the retinal nerve fiber layer (RNFL), ganglion cell layer (GCL), and inner plexiform layer (IPL) which contain retinal ganglion cell axons, their cell bodies, and their synapses with bipolar cells. We did not detect any difference in RNFL, GCL, and IPL combined layer thickness measurements at 500 μm from the optic nerve head. Although, at 1000 μm from the optic nerve head, we found increased thickness in the 9mo C57 mice (12mo C57 p=0.141, 9mo 5xFAD p=0.0005, 12mo 5xFAD p=0.0010) (Fig. 3.C). The lack of outer retinal thinning suggests that structural alterations of the retina such as loss of photoreceptors are likely not responsible for the retinal dysfunction seen in ERG recordings.

**Figure 3.**
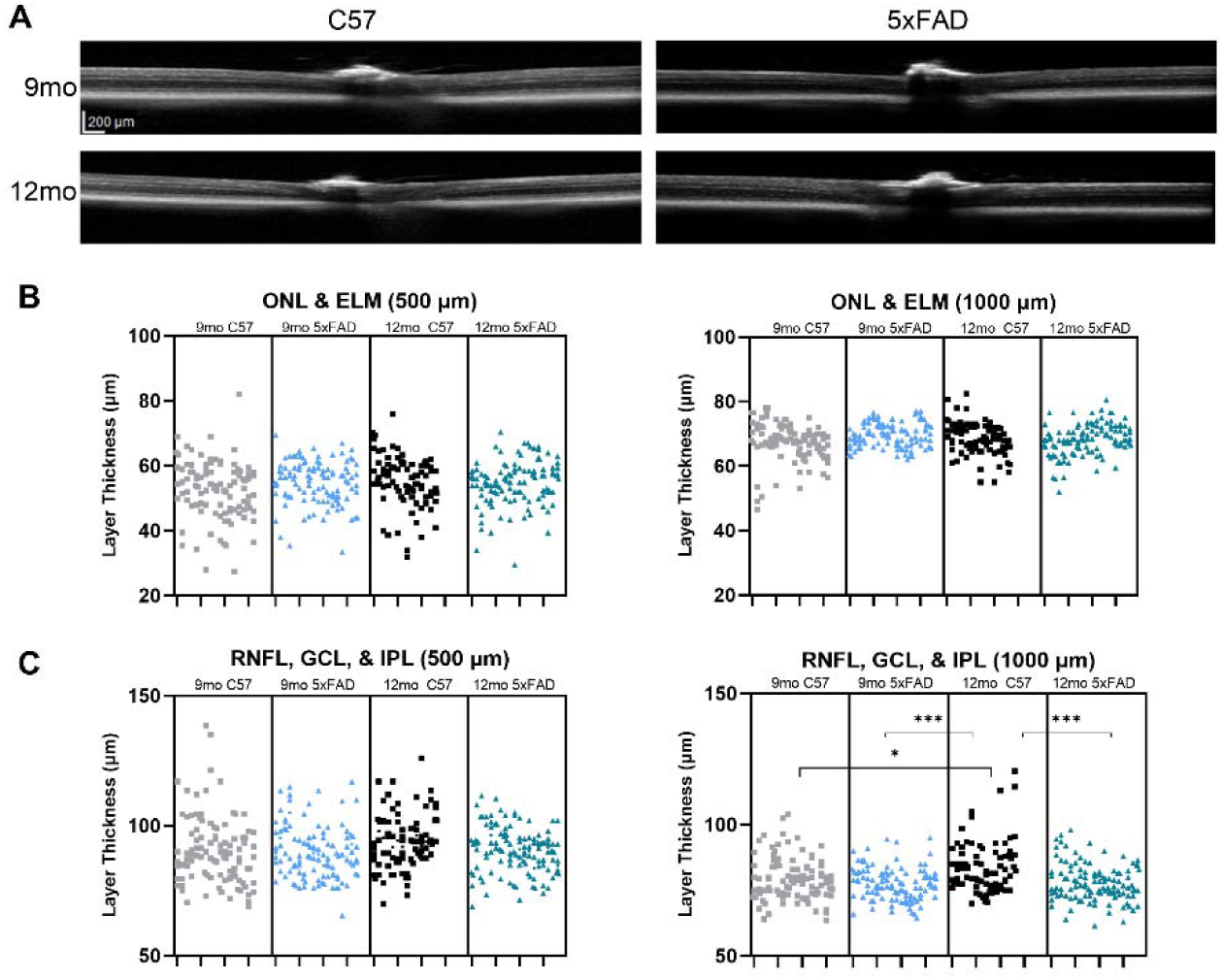
Optical coherence tomography (OCT) of 9- and 12-month mouse retina shows normal retinal structure in the 5xFAD mice. (A) OCT images from 9- and 12-month mice show similar retinal layer thickness in both the C57 and 5xFAD mice. (B) Nested one-way ANOVA of the outer nuclear layer (ONL) and external limiting membrane (ELM) thickness at 500 and 1000 μm from the optic nerve head shows normal layer thickness in the 5xFAD mice. Layer thickness measurements of the retinal nerve fiber layer (RNFL), ganglion cell layer (GCL), and inner plexiform layer (IPL) thickness at 500 and 1000 μm from the optic nerve head show normal retinal structure in the 6- and 9-month 5xFAD mice. Sample size described as number of eyes: n = 17 (9mo C57), 18 (9mo 5xFAD), 14 (12mo C57), 19 (12mo 5xFAD).

We next sought to determine whether amyloid pathology in the 5xFAD mouse leads to dysfunction in a major retinal projection target for conscious vision, the dorsolateral geniculate nucleus (dLGN). To test for the hallmark pathology of AD in the dLGN, we first stained brain sections containing dLGN using thioflavin S to quantify Aβ plaque density. There was widespread pathology in this model in the dLGN as well as in disease-relevant regions such as the hippocampus (Fig. 4.A). We found a high density of Aβ plaques in our region of interest compared to controls which presented with no plaques (Fig. 4.B) verifying our model’s pathology (p<0.0001).

**Figure 4.**
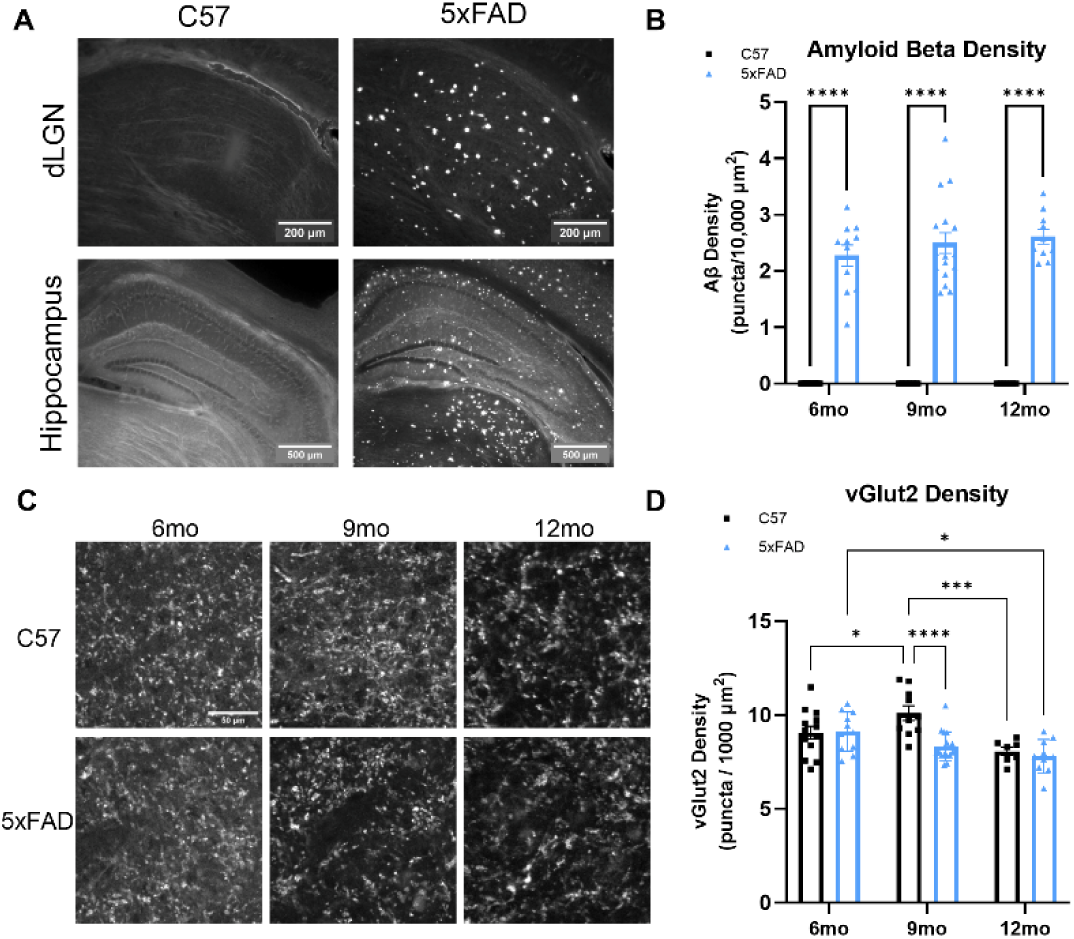
vGlut2 staining of 6-, 9-, and 12-month dLGN sections show accelerated loss of retinal ganglion cell axon terminals in the 5xFAD mice. (A) Example images of C57 and 5xFAD brain sections show Aβ pathology in the dLGN and widespread pathology in a disease-relevant region, the hippocampus. (B) Aβ plaque density was quantified to validate the 5xFAD mouse model. (C) Example images of C57 and 5xFAD brain sections at 3 timepoints show vGlut2 puncta present in the dLGN. (D) Quantification of vGlut2 density in the dLGN shows significant loss of vGlut2 in the 9-month 5xFAD mice compared to controls analyzed by two-way ANOVA with Tukey’s multiple comparisons. Sample size described as number of mice: n = 13 (6mo C57), 11 (6mo 5xFAD), 10 (9mo C57), 17 (9mo 5xFAD), 7 (12mo C57), 10 (12mo 5xFAD).

Next, we investigated the presynaptic structural components of the dLGN, primarily composed of RGC axon terminals. Using 50-μm-thick tissue sections, we stained for vesicular glutamate transporter 2 (vGlut2) to mark RGC axon terminals, and 2-photon images were analyzed for vGlut2 density in the dLGN (Fig. 4C). We found decreased vGlut2 density in the 9mo 5xFAD mice (p<0.0001) with no difference in vGlut2 density between 12mo 5xFAD and control mice again repeating the trend we noted in the retina (Fig. 4D). Following the general pattern we observed in ERG recordings, we found a decrease in RGC axon terminal presence in the 9mo 5xFAD mice compared to age-matched controls, although by 12mo, vGlut2 density was reduced in both mouse lines indicating that Aβ pathology may accelerate an age-dependent loss of RGC axon terminals.

Thalamocortical (TC) neurons are the excitatory neurons of the dLGN and are responsible for relaying information from the retina to higher cortical areas such as the primary visual cortex. With a high presence of Aβ in the dLGN, we next tested whether amyloid pathology is associated with loss of TC neuron somata. To label TC neurons, we stained 50-μm-thick tissue sections for neurons and measured the density of neurons in our tissue sections by counting NeuN+ cells (Fig. 5.A and 5B). Interestingly, we found a significant decrease in cell body density at all three timepoints in the dLGN of 5xFAD mice compared to age-matched controls (6mo p=0.0016, 9mo p=0.0356, 12mo p=0.0003). These data point to pathological diminishment of retinal inputs and degeneration of postsynaptic dLGN neurons along distinct time courses in 5xFAD mice.

**Figure 5.**
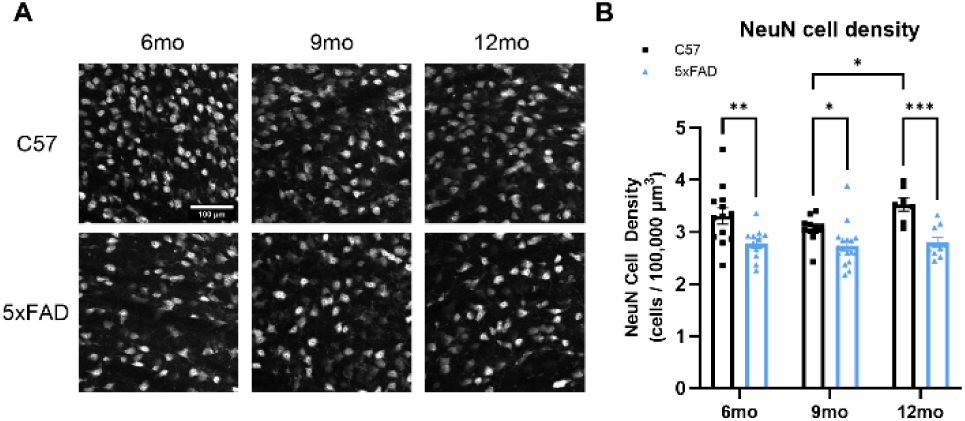
NeuN staining of 6-, 9-, and 12-month dLGN sections shows loss of thalamocortical cell bodies in the 5xFAD mice. (A) Example images of NeuN+ cells in C57 and 5xFAD tissue sections show nuclei counted in the dLGN. (B) Quantification of NeuN+ cells in the dLGN shows significant loss of NeuN at all three timepoints in the 5xFAD mice compared to controls analyzed by two-way ANOVA with Tukey’s multiple comparisons. Sample size described as number of mice: n = 13 (6mo C57), 11 (6mo 5xFAD), 10 (9mo C57), 17 (9mo 5xFAD), 7 (12mo C57), 10 (12mo 5xFAD).

Loss of synapses is a fundamental pathology of AD and is associated with the severity of the disease – both in pathological progression, such as Aβ deposition, and cognitive decline. With this in mind, we performed single neuron dye fills to individually trace TC neuron morphology to determine dendritic wiring in the dLGN (Fig. 6.A and 6.B). After tracing neurons from the dLGN, we analyzed dendritic morphology via Sholl analysis to determine the number of dendrites at increasing radii from the center of the neuron (Fig. 6.C). We then plotted the peak number of intersections of each neuron. Even with the high density of Aβ plaques in the dLGN, the 5xFAD mice did not show alterations in dendritic complexity (Fig. 6.D). Likewise, when we integrated the area under the Sholl curve for each filled TC neuron, there were no detectable signs of dendritic pruning. These results indicate that Aβ does not lead to detectable reorganization of dendrites in the dLGN of the 5xFAD mice up to 9mo of age.

**Figure 6.**
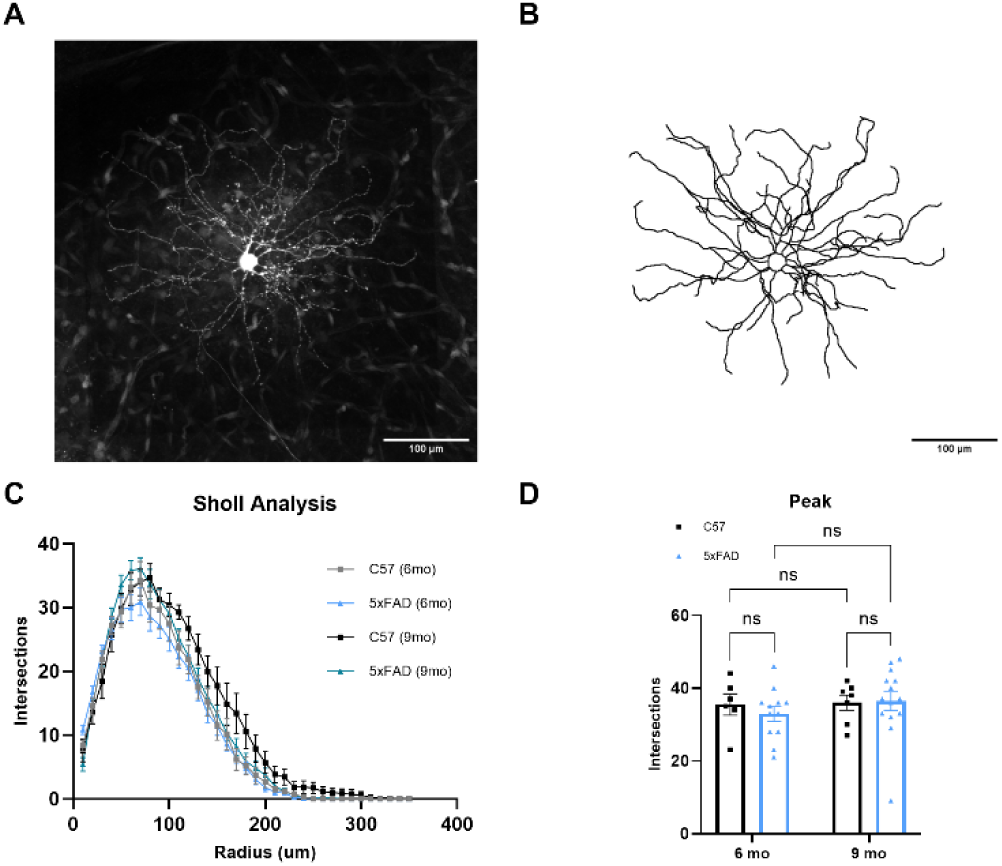
Thalamocortical neuron dendritic structure remains unaltered by Aβ plaques in 5xFAD mice. (A) TC neuron filled with dye shows the cell body and dendrites after 2-photon imaging. (B) An image of the 3D reconstruction of the neuron in ‘A’ shows the same soma and dendrites analyzed by Sholl analysis. (C) Sholl analysis of TC neuron dendritic structure shows normal dendritic composition in the 5xFAD mice. (D) Quantification of peak number of intersections from the Sholl analysis shows no difference in peak amplitude between groups analyzed by two-way ANOVA with uncorrected Fisher’s LSD test. Sample size described as number of cells: n = 6 (6mo C57), 12 (6mo 5xFAD), 7 (9mo C57), 14 (9mo 5xFAD).

Detecting decreased vGlut2 density in the 5xFAD mice indicates there may be alterations in retinogeniculate synaptic function due to loss of RGC axon terminals. To study this, we recorded miniature excitatory postsynaptic currents (mEPSCs) via whole-cell patch-clamp recordings from dLGN in *ex vivo* coronal slices in the absence of stimulation and presence of 60 μM picrotoxin. Amplitude and frequency of events (Fig. 7.B) from individual cells (Fig. 7.A) at 6- and 9-months were analyzed to provide a comparison at time points with detectable vGlut2 loss. We found no change in mEPSC amplitude (Fig. 7.C) when comparing 5xFAD to age-matched control recordings. Interestingly, we did not detect any difference in mEPSC frequency, a parameter of the mEPSCs we would expect to change in response to the decrease in RGC axon terminals in our immunohistochemical analysis (Fig. 7.D). While we have found loss of mEPSC frequency concurrent with vGlut2 loss in a mouse glaucoma model [19,20], we have also found that dLGN mEPSC frequency is unaltered following enucleation surgery [21] which might point to upregulation of remaining RGC synaptic function or possible altered excitatory feedback from the visual cortex.

**Figure 7.**
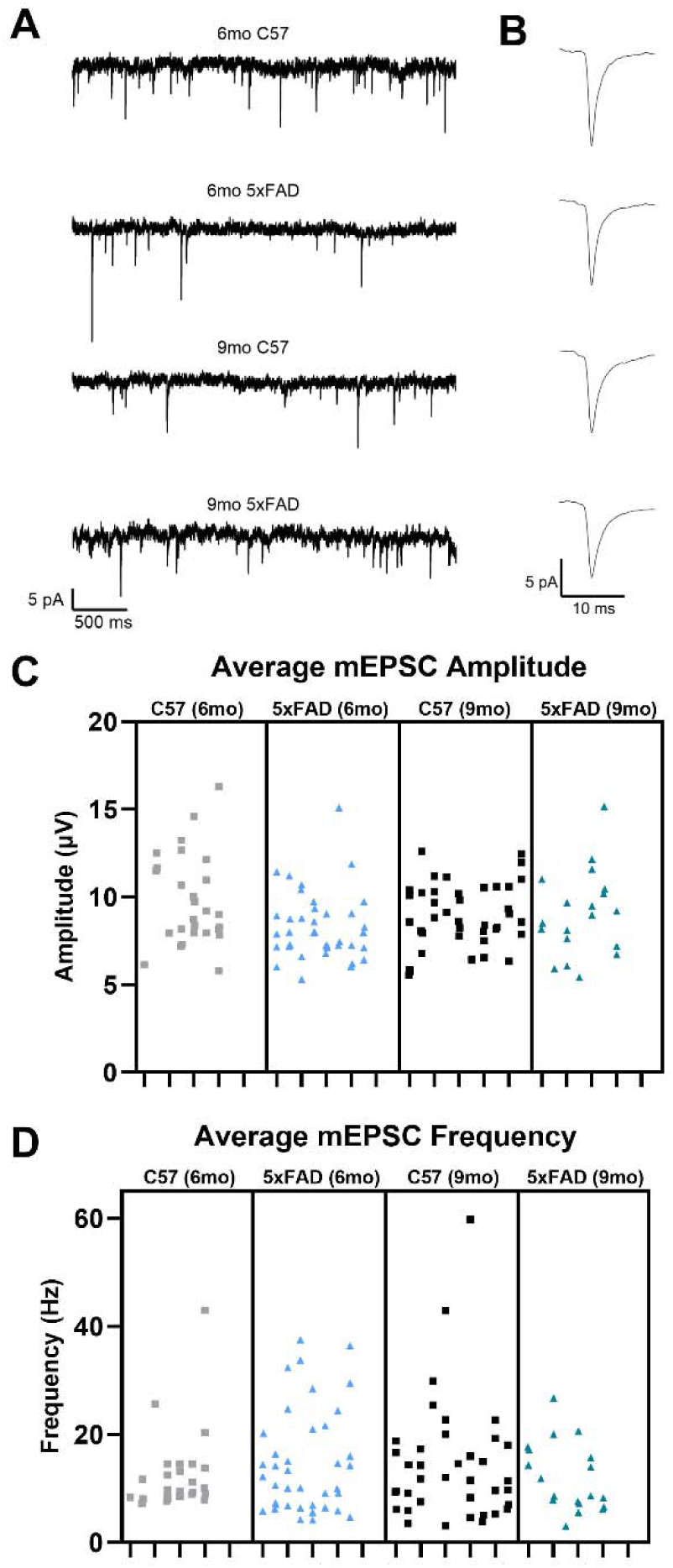
Retinogeniculate synapse function measured via miniature excitatory postsynaptic current (mEPSC) recordings shows normal transsynaptic function. (A) Example traces of miniature excitatory postsynaptic current recordings from TC neurons show amplitude and frequency of events. (B) Example traces of individual events show average event waveform. (C & D) Nested one-way ANOVA with Tukey’s multiple comparisons of mEPSC data shows no change in average mEPSC amplitude (C) or average mEPSC frequency (D) between groups. Sample size described as mice: n = 7 (6mo C57), 8 (6mo 5xFAD), 10 (9mo C57), 7 (9mo 5xFAD). Sample size described as cells: n = 27 (6mo C57), 40 (6mo 5xFAD), 40 (9mo C57), 19 (9mo 5xFAD).

To characterize visual function in the 5xFAD mice from a behavioral perspective, we took advantage of a reflexive visual behavior, optomotor response (OMR). This assesses mouse visual acuity and contrast sensitivity using measures of mouse head movement in response to drifting bar stimuli of varying spatial frequency and contrast. Decreased visual acuity and contrast sensitivity have been reported in patients with both mild and moderate AD [22]. Using vertical sine-wave gratings of differing spatial frequencies (0.025 – 0.45 cpd) or differing contrasts (1 – 96 %), we manually performed a blinded analysis of the percentage of time the mice tracked the stimuli. In our 5xFAD mice, we did not see a decrease in visual acuity or contrast sensitivity indicating normally functioning reflexive vision (Fig. 8.A and 8.B).

**Figure 8.**
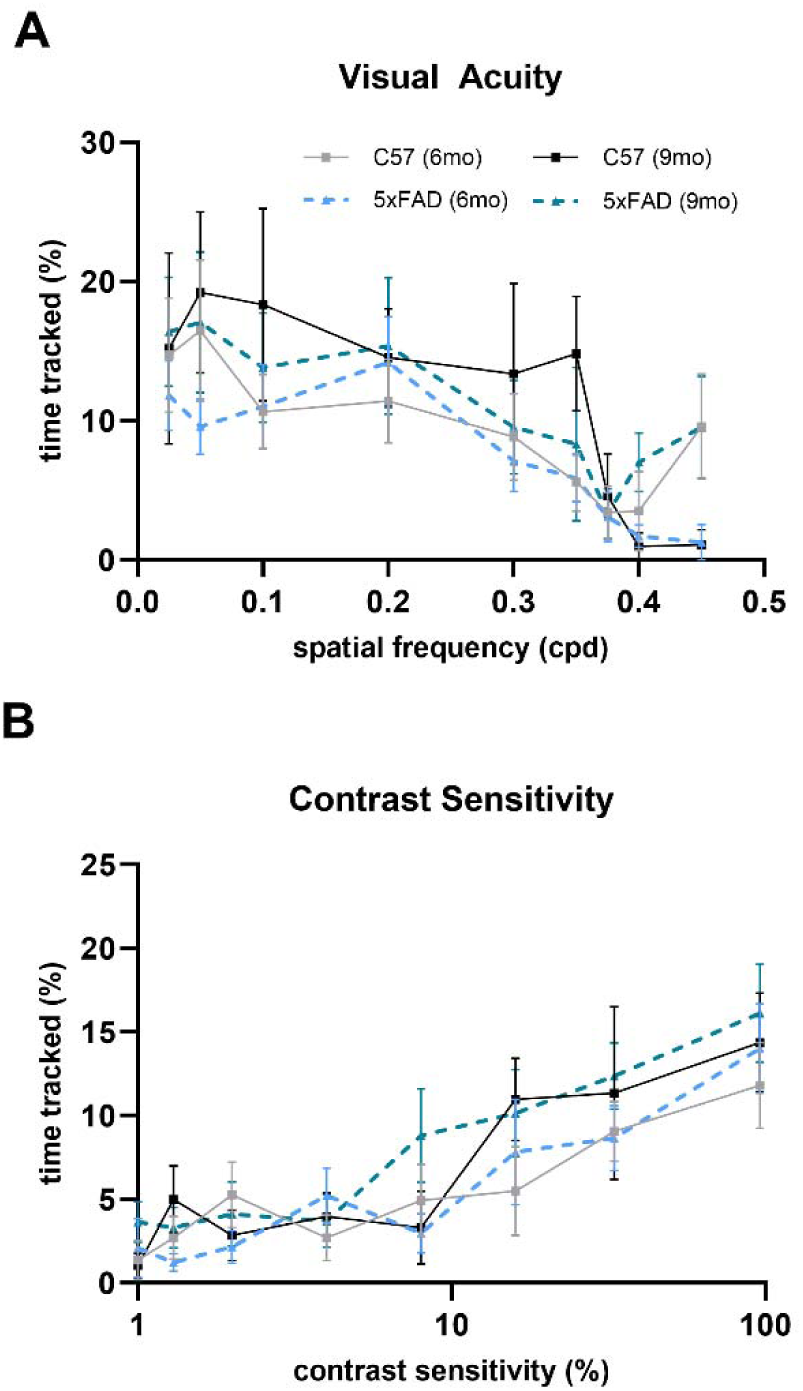
The 5xFAD mouse reflexive visual system functions normally. (A & B) Optomotor response measuring visual acuity (A) and contrast sensitivity (B) shows no change in either parameter in the 5xFAD mice compared to controls analyzed via two-way ANOVA with Tukey’s multiple comparisons. Sample size described as mice: n = 10 (6mo C57), 13 (6mo 5xFAD), 6 (9mo C57), 10 (9mo 5xFAD – visual acuity), 11 (9mo 5xFAD – contrast sensitivity).

## Discussion

A growing number of studies are seeking to link alterations in visual function to neuropathological changes in AD. Our study aims to investigate whether neuronal and synaptic changes that occur in the visual system during AD are due to a specific pathology of the disease, Aβ plaque deposition. This allowed us to parse out the effects of a singular pathology of AD and identify whether these plaques lead to Aβ-induced neurotoxicity causing the visual deficits reported in AD patients or whether they instead might be synergistic effects of Aβ and tau pathology. In this study, we identified functional deficits in the 5xFAD retina and, surprisingly, a lack thereof in the dLGN. Our ERGs displayed altered waveforms indicating photoreceptor impairment in the 6- and 9-month 5xFAD mice, while mEPSC recordings showed normal synaptic function in the dLGN. In terms of structure, retinal layers in the 5xFAD mice had normal thickness as analyzed via OCT, while we found pre- and post-synaptic structural changes in the dLGN including loss of vGlut2-labeled RGC axon terminals and NeuN-labeled neurons in the dLGN through immunohistochemical analysis.

To analyze the retina, the first neural component of the visual pathway, we used clinically relevant, complementary approaches to study its structure and function. With ERG recordings, we concluded there was increased retinal function in the 6mo 5xFAD mice followed by decreased retinal function in the 9mo 5xFAD mice due to altered photoreceptor function. This reversal in pathophysiology may be due to a homeostatic process that preserves visual function in the earlier stages of disease. Altered ERG responses have been noted in this model as well as in other transgenic AD mouse models, but the cause of altered function is unknown [18,23,24]. One explanation particular to the 5xFAD model is disruption by soluble Aβ or Aβ plaques which some have reportedly seen in the retina [18, 25]. Whether the retinal dysfunction is due to Aβ plaques is unclear to us as we were unable to determine whether there are Aβ plaques in the retina of our 5xFAD mice via thioflavin S staining. Due to the lack of substantial evidence that Aβ plaques are forming in the retina of the 5xFAD mouse model, alternate hypotheses require exploration as for the cause of retinal dysfunction. Alternatively, mitochondrial dysfunction in photoreceptors may lead to altered ERG responses. There is evidence of synaptosomal mitochondria dysfunction in the 5xFAD mouse as well as increased mitochondrial oxidative damage seen in 5xFAD whole brain [26,27]. This model was also used to show Aβ pathology-specific dysfunction of mitochondrial metabolism in neurons [28]. In addition to evidence of mitochondrial dysfunction in the 5xFAD mouse, there is also evidence of altered oxidative phosphorylation/mitochondrial pathways in the human AD retina [14]. Photoreceptor bioenergetics of the 5xFAD mouse requires further examination as a potential source of altered ERG responses.

Initially, we considered some structural alteration in the retina may be preceding photoreceptor dysfunction though OCT analysis showed no difference in ONL thickness in the 5xFAD mice. Other reports suggest the 5xFAD model experiences both thinning and thickening of retinal layers via OCT or immunohistochemistry [17,18,25]. In our data set, a 3-month gap between recorded timepoints may be an insufficient time frame to observe structural alterations. Interestingly, we detected an increase in the RNFL, GCL, IPL complex thickness in the 12mo C57 mice. However, we only observed this change at 1000 μm from the optic nerve head. A likely cause of retinal layer thickening via OCT is inflammation in the outer retina of the 12mo C57 mice. This change is statistically significant, but whether it is clinically significant remains to be seen. With ERG and OCT, we show photoreceptor dysfunction in the 5xFAD mice with no change in retinal layer thickness. This suggests that structural alterations to the retina are likely not responsible for the photoreceptor dysfunction.

After passing through the retina, visual information is directed to the dLGN where RGC axons synapse. Synaptic loss has previously been shown in these mice via synaptophysin in whole brain homogenate, and via syntaxin and PSD-95 markers in cortex [29]. In our work, we found a decreased density of vGlut2-labeled RGC axon terminals in the 9mo 5xFAD dLGN. This pathological progression is reminiscent of the decreased retinal function seen in the 9mo 5xFAD ERG recordings.

In addition to the loss of presynaptic RGC axon terminals in the 5xFAD mice, we found decreased neuronal cell body density in the 5xFAD mice in the postsynaptic dLGN at all 3 timepoints. Neuronal cell loss is a pathological event seen in AD following deposition of Aβ plaques and formation of NFTs and has been shown in other brain regions in the 5xFAD mice [30]. As can be seen in this model, neuronal loss still occurs with only the formation of Aβ plaques. Rather than it being an age-dependent phenomenon, it is apparently accelerated in the 5xFAD mice as appears to be the case for vGlut2 loss.

Since there was loss of presynaptic structure in the dLGN, we examined postsynaptic structure by investigating TC neuron dendritic complexity. In AD, rogue synaptic pruning divergent from homeostatic synaptic maintenance takes place and is responsible, at least in part, for the loss of synapses including microglial-mediated synaptic pruning [31]. Knowing this, we examined dendritic wiring of TC neurons in the dLGN. Even with a high burden of Aβ plaques present in the dLGN, an analysis of dendritic complexity (Sholl analysis) showed no change in the dendritic structure of TC neurons in the 5xFAD mouse. Although we see no disruption of dendritic structure in the 5xFAD dLGN, we do see loss of neuronal cell bodies. To reconcile these two findings, we suggest the loss of neuronal cell bodies may not only be due to loss of TC neurons but to loss of interneurons as well. Additionally, aggregation of Aβ in neuronal somata has been shown in this model [29] potentially leading to disruption of somal mitochondria health and somal degeneration preceding dendritic loss.

Considering synaptic function of the 5xFAD mice, deficits in both presynaptic and postsynaptic mechanisms via mEPSC recordings from layer 5 pyramidal neurons have been shown [32]. Altered mEPSC and mIPSC amplitude and frequency have been shown in CA1 neurons of 5xFAD mice [33]. Interestingly, even with fewer RGC axon terminals present in the dLGN, we did not see any changes in mEPSC frequency, a measure of synaptic function we would expect to decrease with fewer retinal inputs when recording from TC neurons. Although this is contradictory to what we expect, we have observed divergent effects on mEPSC frequency previously with reduction in mEPSC frequency in mice with glaucoma and no detectable effects following total loss of retinal inputs resulting from optic nerve transection [19-21]. It is possible that variability makes detection of reduced synaptic function challenging. Although, increases in presynaptic release probability at remaining synapses or upregulation of excitatory corticothalamic feedback may instead be regulating the system to preserve function [34].

Previously, a reflexive visual task known as the optomotor response has shown visual acuity of the 5xFAD mice to be impaired at 6mo [25]. Although the optomotor response is not an output of the conscious visual pathway and is instead a reflexive behavior of the accessory optic system (AOC), we can still extract information applicable to visual function as the AOC receives direct visual input from the retina. With our final piece of data, we found that there was no observable difference in visual acuity or contrast sensitivity of the 5xFAD mice at either the 6mo or 9mo timepoint following a blinded analysis. Decreases in contrast sensitivity and visual acuity are seen in AD patients, but a possible explanation is that other comorbidities of AD such as cataracts, glaucoma, diabetic retinopathy, or age-related macular degeneration are primarily responsible for these visual deficits.

In summary, we found functional deficits in the retina of the 5xFAD mice resulting from altered photoreceptor function. However, we did not detect significant deficits in the function of the dLGN despite RGC synapse loss, neuronal loss, and an overwhelming presence of Aβ plaques in the dLGN. Throughout the dLGN, postsynaptic structural components of the neurons remain intact, and function is maintained. This may be a homeostatic attempt by the visual system to preserve function in the face of pathology. Additionally, it might suggest that other pathological features of AD such as NFTs may play a larger role than Aβ in visual deficits in AD patients.

## Acknowledgements

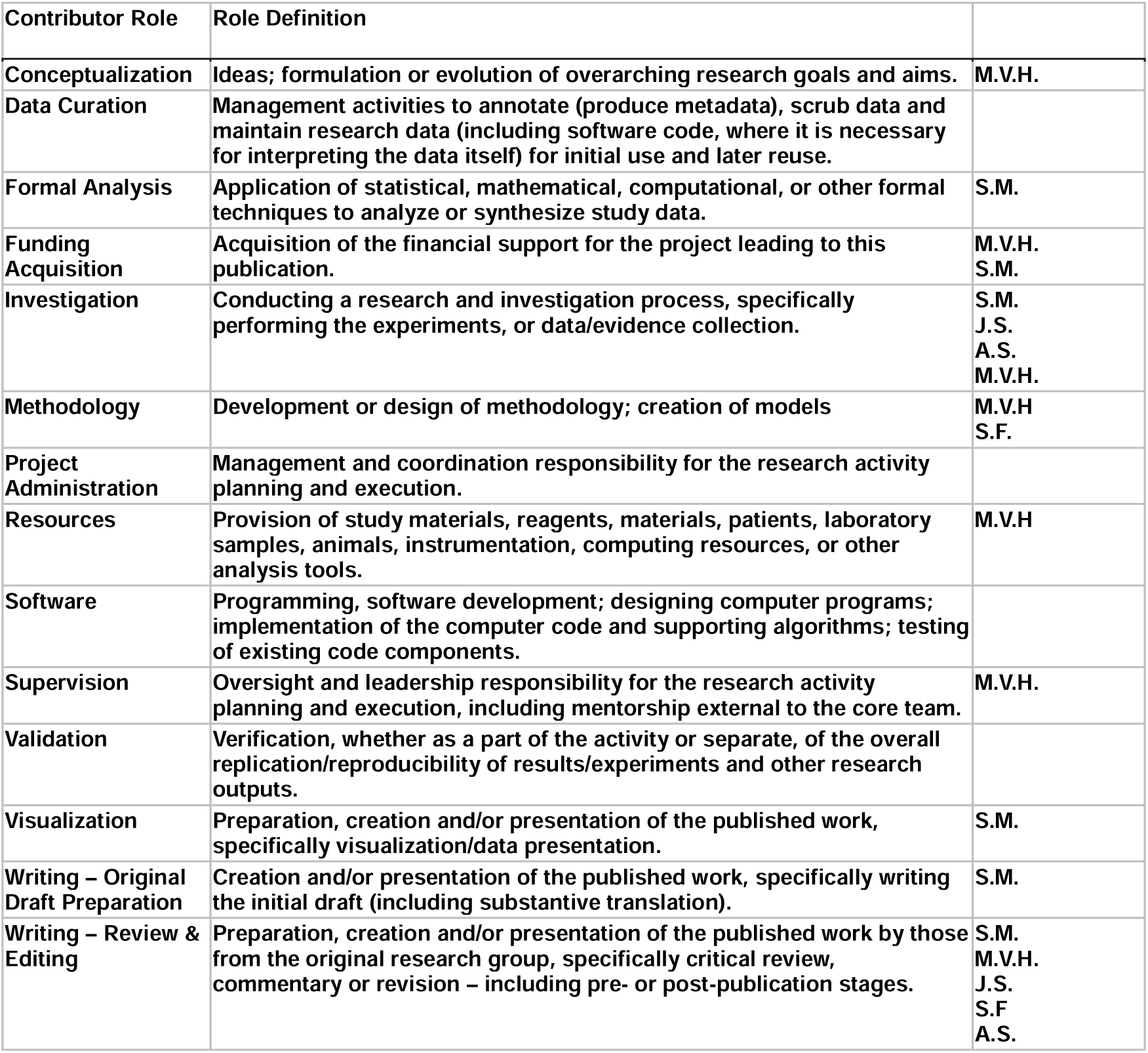

## References

1. Sperling RA, Aisen PS, Beckett LA, Bennett DA, Craft S, Fagan AM, Iwatsubo T, Jack CR Jr, Kaye J, Montine TJ, Park DC, Reiman EM, Rowe CC, Siemers E, Stern Y, Yaffe K, Carrillo MC, Thies B, Morrison-Bogorad M, Wagster MV, Phelps CH. Toward defining the preclinical stages of Alzheimer’s disease: recommendations from the National Institute on Aging-Alzheimer’s Association workgroups on diagnostic guidelines for Alzheimer’s disease. Alzheimers Dement. 2011 May;7(3):280–92. doi: 10.1016/j.jalz.2011.03.003. Epub 2011 Apr 21. PMID: 21514248; PMCID: PMC3220946.

2. A Armstrong R. Risk factors for Alzheimer’s disease. Folia Neuropathol. 2019;57(2):87–105. doi: 10.5114/fn.2019.85929. PMID: 31556570.

3. Selkoe DJ, Hardy J. The amyloid hypothesis of Alzheimer’s disease at 25 years. EMBO Mol Med. 2016 Jun 1;8(6):595–608. doi: 10.15252/emmm.201606210. PMID: 27025652; PMCID: PMC4888851.

4. Vu TA, Fenwick EK, Gan ATL, Man REK, Tan BKJ, Gupta P, Ho KC, Reyes-Ortiz CA, Trompet S, Gussekloo J, O’Brien JM, Mueller-Schotte S, Wong TY, Tham YC, Cheng CY, Lee ATC, Rait G, Swenor BK, Varadaraj V, Brenowitz WD, Medeiros FA, Naël V, Narasimhalu K, Chen CLH, Lamoureux EL. The Bidirectional Relationship between Vision and Cognition: A Systematic Review and Meta-analysis. Ophthalmology. 2021 Jul;128(7):981–992. doi: 10.1016/j.ophtha.2020.12.010. Epub 2021 Feb 27. PMID: 33333104.

5. Ehrlich, J. R., Goldstein, J., Swenor, B. K., Whitson, H., Langa, K. M., & Veliz, P. (2022). Addition of Vision Impairment to a Life-Course Model of Potentially Modifiable Dementia Risk Factors in the US. JAMA Neurology, 79(6), 623–626. 10.1001/jamaneurol.2022.0723

6. Killeen OJ, Zhou Y, Ehrlich JR. Objectively Measured Visual Impairment and Dementia Prevalence in Older Adults in the US. JAMA Ophthalmol. 2023 Aug 1;141(8):786–790. doi: 10.1001/jamaophthalmol.2023.2854. PMID: 37440238; PMCID: PMC10346499.

7. Livingston G, Huntley J, Liu KY, Costafreda SG, Selbæk G, Alladi S, Ames D, Banerjee S, Burns A, Brayne C, Fox NC, Ferri CP, Gitlin LN, Howard R, Kales HC, Kivimäki M, Larson EB, Nakasujja N, Rockwood K, Samus Q, Shirai K, Singh-Manoux A, Schneider LS, Walsh S, Yao Y, Sommerlad A, Mukadam N. Dementia prevention, intervention, and care: 2024 report of the Lancet standing Commission. Lancet. 2024 Aug 10;404(10452):572-628. doi: 10.1016/S0140-6736(24)01296-0. Epub 2024 Jul 31. PMID: 39096926.

8. Kusne Y, Wolf AB, Townley K, Conway M, Peyman GA. Visual system manifestations of Alzheimer’s disease. Acta Ophthalmol. 2017 Dec;95(8):e668–e676. doi: 10.1111/aos.13319. Epub 2016 Nov 19. PMID: 27864881.

9. Javaid FZ, Brenton J, Guo L, Cordeiro MF. Visual and Ocular Manifestations of Alzheimer’s Disease and Their Use as Biomarkers for Diagnosis and Progression. Front Neurol. 2016 Apr 19;7:55. doi: 10.3389/fneur.2016.00055. PMID: 27148157; PMCID: PMC4836138.

10. Ting JT, Daigle TL, Chen Q, Feng G. Acute brain slice methods for adult and aging animals: application of targeted patch clamp analysis and optogenetics. Methods Mol Biol. 2014;1183:221–42. doi: 10.1007/978-1-4939-1096-0_14. PMID: 25023312; PMCID: PMC4219416.

11. Ting JT, Lee BR, Chong P, Soler-Llavina G, Cobbs C, Koch C, Zeng H, Lein E. Preparation of Acute Brain Slices Using an Optimized N-Methyl-D-glucamine Protective Recovery Method. J Vis Exp. 2018 Feb 26;(132):53825. doi: 10.3791/53825. PMID: 29553547; PMCID: PMC5931343.

12. Ferdous S, Liao KL, Gefke ID, Summers VR, Wu W, Donaldson KJ, Kim YK, Sellers JT, Dixon JA, Shelton DA, Markand S, Kim SM, Zhang N, Boatright JH, Nickerson JM. Age-Related Retinal Changes in Wild-Type C57BL/6J Mice Between 2 and 32 Months. Invest Ophthalmol Vis Sci. 2021 Jun 1;62(7):9. doi: 10.1167/iovs.62.7.9. PMID: 34100889; PMCID: PMC8196434.

13. Park JC, Persidina O, Balasubramanian G, Nguyen T, Pradeep A, Hetling JR, McAnany JJ. Effects of normal aging on the mouse retina assessed by full-field flash and flicker electroretinography. Sci Rep. 2023 May 31;13(1):8860. doi: 10.1038/s41598-023-35996-7. PMID: 37258636; PMCID: PMC10232421.

14. Koronyo Y, Rentsendorj A, Mirzaei N, Regis GC, Sheyn J, Shi H, Barron E, Cook-Wiens G, Rodriguez AR, Medeiros R, Paulo JA, Gupta VB, Kramerov AA, Ljubimov AV, Van Eyk JE, Graham SL, Gupta VK, Ringman JM, Hinton DR, Miller CA, Black KL, Cattaneo A, Meli G, Mirzaei M, Fuchs DT, Koronyo-Hamaoui M. Retinal pathological features and proteome signatures of Alzheimer’s disease. Acta Neuropathol. 2023 Apr;145(4):409–438. doi: 10.1007/s00401-023-02548-2. Epub 2023 Feb 11. PMID: 36773106; PMCID: PMC10020290.

15. Wachtmeister L, Dowling JE. The oscillatory potentials of the mudpuppy retina. Invest Ophthalmol Vis Sci. 1978 Dec;17(12):1176–88. PMID: 721390.

16. Liao F, Liu H, Milla-Navarro S, Villa P, Germain F. Origin of Retinal Oscillatory Potentials in the Mouse, a Tool to Specifically Locate Retinal Damage. Int J Mol Sci. 2023 Feb 4;24(4):3126. doi: 10.3390/ijms24043126. PMID: 36834538; PMCID: PMC9958948.

17. Kim TH, Son T, Klatt D, Yao X. Concurrent OCT and OCT angiography of retinal neurovascular degeneration in the 5XFAD Alzheimer’s disease mice. Neurophotonics. 2021 Jul;8(3):035002. doi: 10.1117/1.NPh.8.3.035002. Epub 2021 Jul 10. PMID: 34277888; PMCID: PMC8271351.

18. Lim JKH, Li QX, He Z, Vingrys AJ, Chinnery HR, Mullen J, Bui BV, Nguyen CTO. Retinal Functional and Structural Changes in the 5xFAD Mouse Model of Alzheimer’s Disease. Front Neurosci. 2020 Aug 13;14:862. doi: 10.3389/fnins.2020.00862. PMID: 32903645; PMCID: PMC7438734.

19. Smith JC, Zhang KY, Sladek A, Thompson J, Bierlein ER, Bhandari A, Van Hook MJ. Loss of Retinogeniculate Synaptic Function in the DBA/2J Mouse Model of Glaucoma. eNeuro. 2022 Dec 27;9(6):ENEURO.0421-22.2022. doi: 10.1523/ENEURO.0421-22.2022. PMID: 36526366; PMCID: PMC9794376.

20. Van Hook MJ, Monaco C, Bierlein ER, Smith JC. Neuronal and Synaptic Plasticity in the Visual Thalamus in Mouse Models of Glaucoma. Front Cell Neurosci. 2021 Jan 15;14:626056. doi: 10.3389/fncel.2020.626056. PMID: 33584206; PMCID: PMC7873902.

21. Bhandari A, Ward TW, Smith J, Van Hook MJ. Structural and Functional Plasticity in the Dorsolateral Geniculate Nucleus of Mice following Bilateral Enucleation. Neuroscience. 2022 Apr 15;488:44–59. doi: 10.1016/j.neuroscience.2022.01.029. Epub 2022 Feb 4. PMID: 35131394; PMCID: PMC8960354.

22. Salobrar-García E, de Hoz R, Ramírez AI, López-Cuenca I, Rojas P, Vazirani R, Amarante C, Yubero R, Gil P, Pinazo-Durán MD, Salazar JJ, Ramírez JM. Changes in visual function and retinal structure in the progression of Alzheimer’s disease. PLoS One. 2019 Aug 15;14(8):e0220535. doi: 10.1371/journal.pone.0220535. PMID: 31415594; PMCID: PMC6695171.

23. Perez SE, Lumayag S, Kovacs B, Mufson EJ, Xu S. Beta-amyloid deposition and functional impairment in the retina of the APPswe/PS1DeltaE9 transgenic mouse model of Alzheimer’s disease. Invest Ophthalmol Vis Sci. 2009 Feb;50(2):793–800. doi: 10.1167/iovs.08-2384. Epub 2008 Sep 12. PMID: 18791173; PMCID: PMC3697019.

24. McAnany JJ, Matei N, Chen YF, Liu K, Park JC, Shahidi M. Rod pathway and cone pathway retinal dysfunction in the 5xFAD mouse model of Alzheimer’s disease. Sci Rep. 2021 Mar 1;11(1):4824. doi: 10.1038/s41598-021-84318-2. PMID: 33649406; PMCID: PMC7921657.

25. Zhang M, Zhong L, Han X, Xiong G, Xu D, Zhang S, Cheng H, Chiu K, Xu Y. Brain and Retinal Abnormalities in the 5xFAD Mouse Model of Alzheimer’s Disease at Early Stages. Front Neurosci. 2021 Jul 23;15:681831. doi: 10.3389/fnins.2021.681831. PMID: 34366774; PMCID: PMC8343228.

26. Wang L, Guo L, Lu L, Sun H, Shao M, Beck SJ, Li L, Ramachandran J, Du Y, Du H. Synaptosomal Mitochondrial Dysfunction in 5xFAD Mouse Model of Alzheimer’s Disease. PLoS One. 2016 Mar 4;11(3):e0150441. doi: 10.1371/journal.pone.0150441. PMID: 26942905; PMCID: PMC4778903.

27. Yoshida N, Kato Y, Takatsu H, Fukui K. Relationship between Cognitive Dysfunction and Age-Related Variability in Oxidative Markers in Isolated Mitochondria of Alzheimer’s Disease Transgenic Mouse Brains. Biomedicines. 2022 Jan 26;10(2):281. doi: 10.3390/biomedicines10020281. PMID: 35203488; PMCID: PMC8869326.

28. Kuhn MK, Fleeman RM, Beidler LM, Snyder AM, Chan DC, Proctor EA. Amyloid-β Pathology-Specific Cytokine Secretion Suppresses Neuronal Mitochondrial Metabolism. Cell Mol Bioeng. 2023 Sep 11;16(4):405–421. doi: 10.1007/s12195-023-00782-y. PMID: 37811007; PMCID: PMC10550897.

29. Oakley H, Cole SL, Logan S, Maus E, Shao P, Craft J, Guillozet-Bongaarts A, Ohno M, Disterhoft J, Van Eldik L, Berry R, Vassar R. Intraneuronal beta-amyloid aggregates, neurodegeneration, and neuron loss in transgenic mice with five familial Alzheimer’s disease mutations: potential factors in amyloid plaque formation. J Neurosci. 2006 Oct 4;26(40):10129–40. doi: 10.1523/JNEUROSCI.1202-06.2006. PMID: 17021169; PMCID: PMC6674618.

30. Eimer WA, Vassar R. Neuron loss in the 5XFAD mouse model of Alzheimer’s disease correlates with intraneuronal Aβ42 accumulation and Caspase-3 activation. Mol Neurodegener. 2013 Jan 14;8:2. doi: 10.1186/1750-1326-8-2. PMID: 23316765; PMCID: PMC3552866.

31. Hong S, Beja-Glasser VF, Nfonoyim BM, Frouin A, Li S, Ramakrishnan S, Merry KM, Shi Q, Rosenthal A, Barres BA, Lemere CA, Selkoe DJ, Stevens B. Complement and microglia mediate early synapse loss in Alzheimer mouse models. Science. 2016 May 6;352(6286):712-716. doi: 10.1126/science.aad8373. Epub 2016 Mar 31. PMID: 27033548; PMCID: PMC5094372.

32. Buskila Y, Crowe SE, Ellis-Davies GC. Synaptic deficits in layer 5 neurons precede overt structural decay in 5xFAD mice. Neuroscience. 2013 Dec 19;254:152–9. doi: 10.1016/j.neuroscience.2013.09.016. Epub 2013 Sep 20. PMID: 24055684; PMCID: PMC4078998.

33. Barbour AJ, Gourmaud S, Lancaster E, Li X, Stewart DA, Hoag KF, Irwin DJ, Talos DM, Jensen FE. Seizures exacerbate excitatory: inhibitory imbalance in Alzheimer’s disease and 5XFAD mice. Brain. 2024 Jun 3;147(6):2169–2184. doi: 10.1093/brain/awae126. PMID: 38662500; PMCID: PMC11146435.

34. Krahe TE, Guido W. Homeostatic plasticity in the visual thalamus by monocular deprivation. J Neurosci. 2011 May 4;31(18):6842–9. doi: 10.1523/JNEUROSCI.1173-11.2011. PMID: 21543614; PMCID: PMC3319043.

